# Contrastive modelling of transcription and transcript abundance in legumes using PlanTT

**DOI:** 10.1101/2025.11.15.685414

**Authors:** Nicolas Raymond, Xi Zhang, Jubair Sheikh, Fatima Daveouis, Ruchika Verma, Dustin Cram, Halim Song, Yongguo Cao, Morgan Kirzinger, Deborah Akaniru, Jordan Ubbens, David Konkin

## Abstract

Predicting the impacts of sequence variation on gene expression remains a challenging task. Further, in plants, we have a limited understanding of the relative contributions of different gene expression regulatory mechanisms. To address these limitations we generated a comparative multiomic dataset comprising matched 3’-RNA-seq and PRO-seq data from matched tissues of reference genotypes of four legumes of the invert repeat lacking clade (*Pisum sativum, Vicia faba, Lathyrus sativa* and *Medicago truncatula*). Focused on the challenging task of predicting expression differences between ortholog pairs from unseen orthogroups, we used this dataset and a novel prediction framework to build contrastive models that predict quantitative differences (effect size differences) in transcription and transcript abundance.

## 1. Introduction

Although climate change has potential to temporarily increase crop yield in certain regions due to extended growing seasons [1], food security is expected to be compromised by more frequent extreme weather events (e.g., flooding, drought) [2]. Increases in crop productivity and stability are needed to tackle food insecurity driven by population growth and climate factors. One opportunity to accelerate crop development is via integration of knowledge of intermediate biological processes such as gene expression into crop development strategies.

Gene expression is the process by which a gene is converted to a functional product. Genes encoded in DNA are first transcribed to RNA transcripts and, for protein-coding genes, a subclass of RNAs called messenger RNAs (mRNAs) are translated into proteins. Spatiotemporal patterns and responsiveness of gene expression plays a critical role in organisms development and homeostasis. Therefore, modelling how the information encoded in DNA regulates transcript abundance is central to understanding the impacts of DNA sequence variation on an organism’s phenotype.

Modern sequencing technologies allow both determination of the genetic sequence and global measurement of mRNA abundance [3, 4]. Further, deep learning models such as convolutional neural networks (CNNs) and transformers shown promise for modelling the relationship between nucleotide sequences and gene expression [5, 6, 7]. However, many of the efforts to model gene expression as a function of sequence focus on developing models capable of predicting the expression of different genes within the genome of a single individual. While these models demonstrate impressive ability to reconstitute gene expression tracks and predict expression differences across the different genes within an individual, they fail at the task of predicting expression differences between different alleles of the same gene across different individuals [8]. Given that this task of allelic prediction may be the most important use case for applying these models to crop improvement strategies, including variant interpretation, genomic prediction, and gene editing, advances to modelling approaches are needed.

In order to focus gene expression models on allelic differences, Washburn et al. presented the idea of training models on contrasting pairs of related genes, and applied this method in Maize and Sorghum orthologs to classify which ortholog has higher expression. Adopting a pairwise contrastive approach offers a means to control for evolutionary confounders such as expression patterns associated with classes of genes or protein domains. Stated more plainly, contrastive approaches can control for the type of gene so the model can focus on variation directly impacting expression.

Two challenges that limit the application of contrastive approaches to quantitative gene expression prediction are 1) a lack of high-quality datasets comprised of multiple species collected in parallel to eliminate experimental confounders, and 2) challenges with using log-scaled expression differences as a predictor. To address the first limitation we generated a highly replicated, matched tissue expression dataset for reference accessions of four related legume species field pea (*Pisum sativum*), faba bean (*Vicia faba*), grass pea (*Lathyrus sativus*), and barrel medic (*Medicago truncatula*). With this dataset, we implement a model architecture designed to predict differential gene expression between orthologs in terms of effect size. This mitigates the difficulties of using log-scaled differences and directly accounts for variance in expression measurement.

Gene expression is governed at multiple layers including transcription and transcript degradation. Transcript abundance, as measured by RNA-seq is a function of both of these processes. Methods that capture active transcription, such as precision run on sequencing (PRO-seq), provide a direct measurement of transcription, and when coupled with matched RNA-seq, can provide estimates of transcript stability. We supplement our matched multi-species transcriptomic dataset using PRO-seq revealing that promoter proximal pausing is widespread in vicioid legumes and providing training data for a contrastive model of transcription. Finally, we explore potential synergies of using these data together.

## 2. Theoretical background

### 2.1. Foundations of contrastive models

In this work we first laid the foundations of contrastive models for gene expression difference. Precisely, we established an exhaustive list of mathematical criteria that must be satisfied by a function to be considered as a valid model to predict the difference of expression between two genes based on nucleotide sequences associated to both of them respectively (Definition 1).

#### Definition 1.

*Let* **s**_*a*_, **s**_*b*_ ∈ ℝ^*D×L*^ *be sequences of L nucleotides, for which each nucleotide is encoded using a D-dimensional vector. Let us also consider that* **s**_*a*_ *and* **s**_*b*_ *are associated respectively to genes a and b of unknown expression levels e*_*a*_ *and e*_*b*_. *A function f* : (ℝ^*D×L*^, ℝ^*D×L*^) *→* ℝ *is a valid prediction model of gene expression difference (i.e*., *e*_*a*_ *− e*_*b*_*) if it satisfies the two following criteria:*

1. *f* (**s**_*a*_, **s**_*a*_) = 0
2. *f* (**s**_*a*_, **s**_*b*_) = *−f* (**s**_*b*_, **s**_*a*_)

In Definition 1, the first criterion specifies that the prediction of the model must be null if a sequence is compared to itself. The second indicates that the value predicted by the model should be of the same magnitude, but of the opposite sign, if the input sequences are permuted. Differently put, the model must be *equivariant* to the order of the sequences.

### 2.2. PlanTT

Here we establish the construction of a contrastive model predicting the mRNA abundance difference between two versions of a gene using their corresponding nucleotide sequences, which we refer to as the **T**wo-**T**ower contrast model for **Plan**t genes (PlanTT), (Figure 1). PlanTT has a generic architecture composed of two towers with shared weights, and a head. The towers embed genes’ sequence information into a lower-dimensional representation, and the head uses the difference of these embeddings to resolve a prediction of gene expression difference.

**Figure 1.**
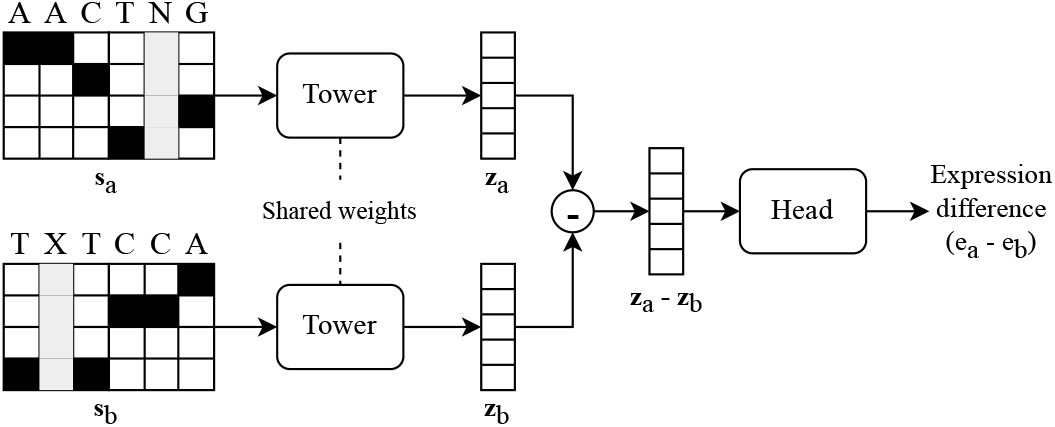
Illustration of the **t**wo-**t**ower contrast model for **plan**t genes (PlanTT). Encoded reference sequence **s**_*a*_ and edited sequence **s**_*b*_ are first fed separately to the towers to generate their respective embedding **z**_*a*_ and **z**_*b*_. The difference of the embeddings (i.e., **z**_*a*_ *−* **z**_*b*_) is further passed to the head of the model to return a prediction of the expression difference between the two genes (i.e., *e*_*a*_ − *e*_*b*_). Parameters of the towers and the head are learned in an end-to-end fashion during the training.

We conclude that using an odd function for the head is a sufficient condition for PlanTT to satisfy Definition 1 (Supplementary Theorem 1). Proof of this theorem is given in Supplementary Proof 1. Therefore, we defined the head of our model as the element-wise sum of the difference of the embedding vectors (i.e., **z**_*a*_ − **z**_*b*_). The odd nature of this head function is proven in Supplementary Proof 2. In addition, this choice of head function is relevant because it can be re-written has the difference between the sum of the elements in each embedding:

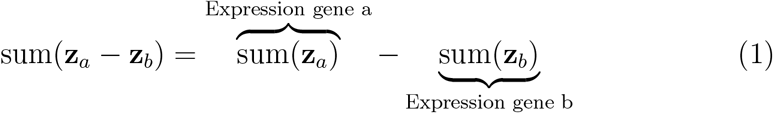

## 3. Materials and methods

### 3.1. Data generation

Reference accessions for *P. sativum, V. faba, M. truncatula* and *L. sativus* were grown together in a controlled environment (18h photoperiod, 22°C day 18°C Night). The basal two leaflets of the third leaf were collected into liquid nitrogen when the fifth leaf had expanded. Six replicates were acquired per species.

For 3’-RNA-seq, RNA was isolated individually from each of the six replicates using a Monarch Total RNA kit (NEB). 3’-RNA-seq libraries were prepared using a QuantSeq [4] 3’ FWD mRNA-Seq Library Prep Kit (Lexogen) and sequenced on an Illumina Novaseq 6000. Reads were trimmed and mapped with STAR [9]. The average counts per million (CPM) of the six replicates was considered as the mRNA abundance for each gene.

Three replicates of Precision Run-On sequencing (PRO-seq) data were generated for each species from the same RNA used for 3’-RNA-seq data by pooling RNA from two plants for each replicate. PRO-seq libraries were prepared according to [10], and sequenced on an Illumina HiSeq instrument with paired 150 bp reads. Reads were trimmed using Cutadapt v3.5 [11] with parameters -O 10 -a TGGAATTCTCGG -A GATCGTCGGACT -q 15 -m 40:40 -j 2. The trimmed reads were then aligned and mapped by STAR v2.7.10 [9] with default parameters. Finally, we normalized read counts by converting them to average counts per million (CPM) and transcripts per million (TPM) [12] based on the length of each transcript. The metagene plot of PRO-seq data was implemented in Deeptools2 [13] via the built-in functions plotHeatmap and plotProfile, and the computeMatrix function was used to capture the signal over contiguous regions around TSS and TTS.

#### 3.1.1. Paired sequence extraction

OrthoFinder v2.5.5 [14] was used to identify orthogroups. Orthologs were used if they were the sole ortholog for a species within the orthogroup and were at least 1kb in length. In total 65,495 orthologs pairs were utilized (Supplementary Figure 1). Each gene in each pair of orthologs was represented using a sequence of 3000 bases, consisting of 1Kb upstream of the gene annotation, the first 500 bases of the annotated gene, the last 500 bases of the annotated gene, and 1Kb downstream of the gene (Figure 2). The nucleotide sequence was one hot-encoded. As many genes in the current annotations of these species are missing UTR annotation, the three nucleotides starting at the TSS and the three ending at the TTS were masked to prevent the model learning from translational start and stop codons.

**Figure 2.**
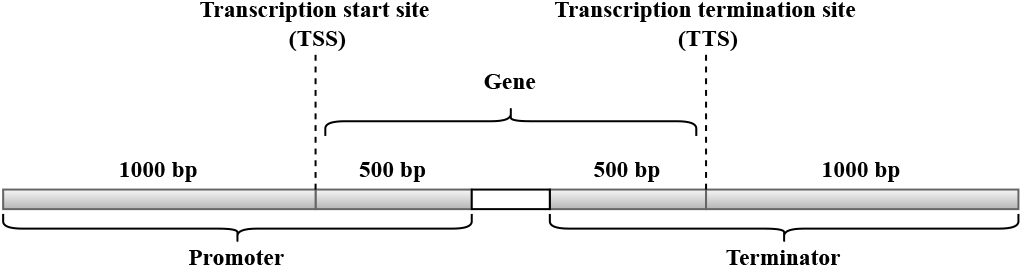
Features associated to a single gene. In green, a sequence of 3000 base pairs (bp) is equally separated around transcription start site (TSS) and transcription termination site (TTS). The figure is inspired from previous work [7].

Given the mRNA abundance readings of a pair of orthologs 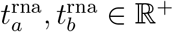, the goal of the proposed contrastive model is to predict the difference in expression between 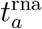 and 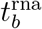. This raises an issue when deciding on the regression target. This regression problem is distinct from previous work which aims to quantify only the magnitude of gene expression for a syntenic single-copy ortholog pairs, or to predict a binary classification to distinguish between 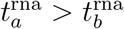 and 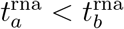 [7]. To be noted, synteny-based single-copy gene pair identification is highly dependent on genome assembly quality [15], which helps mitigate genes involving in tandem duplications or recent gene losses but risks creating non-balanced datasets. Here, the proposed target needs to be capable of capturing both the direction of the difference as well as its magnitude. Predicting the difference in the raw read counts is intractable due to the fact that they are pareto-distributed (since the read counts themselves are exponentially distributed) – however, predicting the log of this difference log 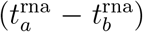 is also unsuitable as this destroys the ability to produce negative predictions in the case where 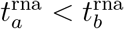. Another option is to predict the “difference of logs” 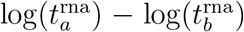. Although this numerically allows for both magnitude and direction, this is equivalent to predicting 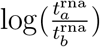 which implies predicting the log of a ratio. This is problematic as ratios explode asymptotically as the denominator approaches zero. Because of very many expression values close to zero, this is not an informative target as the loss signal is dominated by these samples and any pair having significant 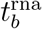 is essentially ignored. It also means that any pairs where either copy is unexpressed must be removed. The targets can be smoothed by adding an arbitrary constant to each side of the ratio, but this means that the network is no longer trained to minimize the difference of logs, and the predictions from the model are no longer in any meaningful unit (and cannot be reverted to one).

In order to predict both the magnitude and direction in a way which reflects the probability that a difference is significant, we predict Cohen’s *d*, a statement of effect size. Since each expression measurement is a mean over multiple replicates, standard deviations *σ* are available for each one. These can be used to compute *d* for an ortholog pair:

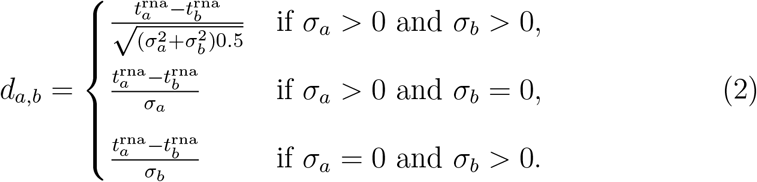

Cohen’s *d* intuitively represents the difference in expression, normalized by the expected variance of the measurements. This allows *d* to represent large and small differences regardless of the absolute magnitude of the expression. It also takes into account the noisy nature of mRNA read counts, as the effect size shrinks with the variance of the measurements, naturally reducing the chances of false positives. Predicting effect size allows us to put the expression differences into context without relying on ratios, which are numerically problematic.

Six and three replicates were used for the 3’-RNA-seq and PRO-seq experiments, respectively. This is a small sample size for computing Cohen’s *d*, which can become noisy at small sample sizes due to unreliable estimates of mean and variance. However, subsequent experiments wherein the measured variance for each gene was replaced with a predicted variance obtained by linear regression on the magnitude of the expression yielded poorer predictive performance despite fewer outliers in the effect size labels. This implies that PlanTT is better at predicting the extremely high effect sizes, which could suggest that these large changes are in fact not artifacts of small sample size but are due to biologically predictable effects. Although we are confident that our effect size estimates are robust, sample size remains a limitation in all experiments aiming to quantify gene expression.

#### 3.1.2. Training, validation and test sets

Although each pair of orthologs was, as a whole, distinct from all the others included in the dataset, it often shared a common gene with another pair. More precisely, this situation occurred when a gene was part of an *orthogroup* with more than two members. An orthogroup is a set of genes that are considered as being single-copy orthologs (i.e., equivalent) across at least two of the four species in the dataset. It can therefore contain up to four members – one gene from each of the species (*M. truncatula, V. faba, P. sativum*, and *L. sativus*). The number of distinct and unordered pairs of genes attached to a single orthogroup of *n* members (i.e. constituents) can be calculated with the binomial coefficient:

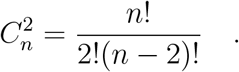

Because test pairs comprised of genes already appearing in the training or validation set could inflate the performance measured during experiments, we conducted our model training and testing using orthogroup-guided crossvalidation splits. We created five stratified cross-validation splits by sampling orthogroups rather than pairs of genes. The composition of each of the training, validation and test sets was stratified in terms of orthogroup labels. The latter labels were established by saving a set with the first letters of each of the species from which the genes in the orthogroup were taken. For example, an orthogroup with members from *M. truncatula* and *V. faba* had the label {*M, V*}. Each training, validation, and test set was further balanced to contain an approximately even number of positive and negative targets. The balancing was performed by permuting the input sequences (**s**_*a*_, **s**_*b*_ *→* **s**_*b*_, **s**_*a*_), and switching the target signs (*t* → *− t*), of certain data points with the least represented target sign.

### 3.2. Tower architecture (PlanTT-CNN)

As presented in Section 2.2, PlanTT is a generic model for which the shared tower architecture needs to be determined. Here, taking inspiration from ResNet [16] we built a deep convolutional architecture that integrates residual blocks (Figure 3).

**Figure 3.**
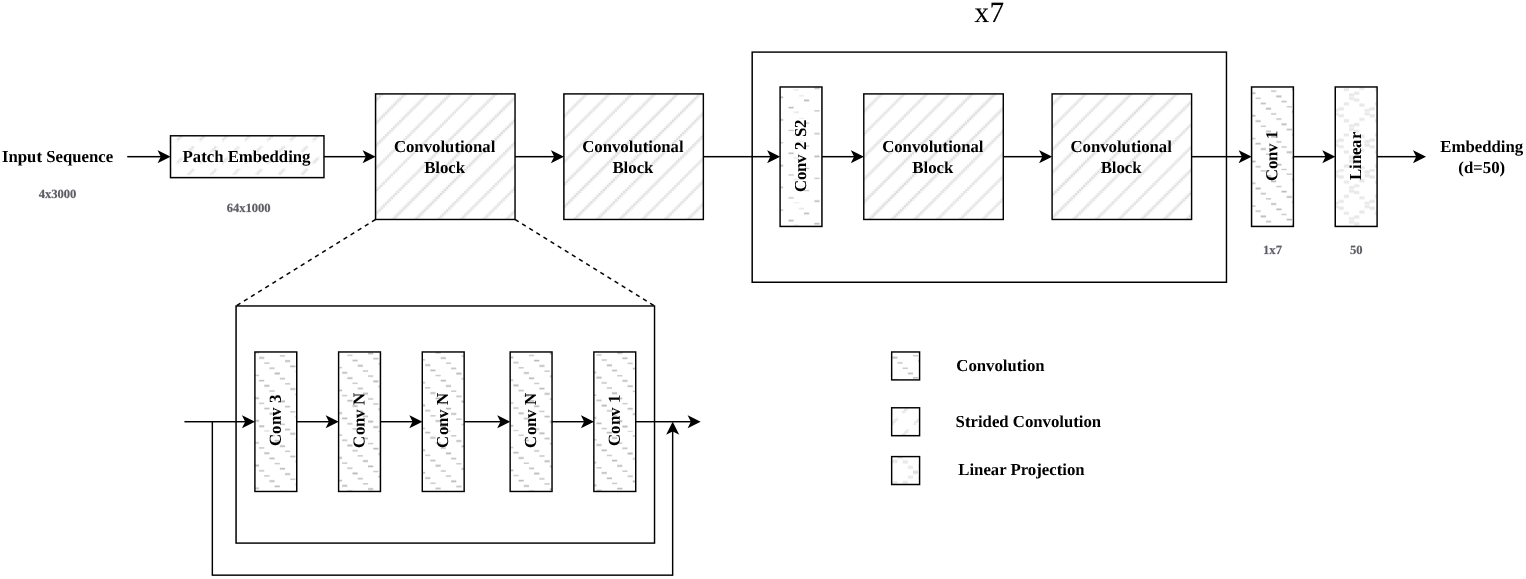
High-level overview of the tower architecture.

For this tower, each nucleotide of the 3kb long sequences was soft encoded using the following probability distributions: *A* = [1, 0, 0, 0], *C* = [0, 1, 0, 0], *G* = [0, 0, 1, 0], *T* = [0, 0, 0, 1] and *N* = *X* = [.25,.25,.25,.25]. The input stem consists of a patch embedding which downsamples the input sequence by a factor of three and outputs a volume with a depth of 64 channels. The network is comprised of 8 stages, each incorporating two convolutional blocks. The latter 7 stages are initiated with a strided convolution which downsamples the input by a factor of two, and doubles the depth up to a maximum depth of 256 channels. The convolutional blocks are equipped with a residual connection and consist of a convolution operation with a receptive field of 3, three convolution operations with a receptive field dependent on the stage, and finally a pointwise convolution. The receptive field for the inner convolutions are 11, 9, 7, 7, 5, 5, 3, and 3 for stages 1, 2, 3, etc. Each convolution operation is followed by batch normalization [17] and dropout [18]. The output from the final convolutional block is followed by a final pointwise convolution with an output depth of 1, and then a linear projection to a 50-dimensional output embedding.

### 3.3. Training settings

For each model training, we employed the AdamW [19] optimizer with parameters *β*_1_ = 0.9, *β*_2_ = 0.999, *ϵ* = 1*e* − 08, and weight decay = 0.01. A single-cycle learning rate schedule was used with a warmup phase of 900 steps, a maximum learning rate of 8*e* − 5, and cosine decay back to zero. All models were trained with a batch size of 32. For both PlanTT and the Washburn et al. model [7], we used early stopping with a budget of 50 epochs and a patience of 10 epochs. For PlanTT, we used the mean squared error (MSE) as the loss function and Spearman’s *r* as the metric for early stopping.

## 4. Results

### 4.1. Matched RNA-seq and PRO-seq analyses support the important role for transcript stability in regulation of transcript abundance in legumes

To investigate gene regulation in inverted repeat-lacking clade (IRLC) legumes and all cross-species comparative transcriptomics at the level of transcript abundance and active transcription, we acquired 3’-RNA-seq and PRO-seq data from developmentally matched leaflet tissues for reference lineages of *Pisum sativum* (cv Cameor), *Vicia faba* (cv Hedin/2), *Lathyrus sativa* (cv LS007) and *Medicago truncatula* (cv Jemalong A17). Plants were grown concurrently in the same controlled environment and ground tissues were split for PRO-seq and 3’-RNA-seq measurements. Both methods displayed high replicate concordance (with average Pearson R^2^ values above 0.8 for both sequencing methods, Supplementary Figures 2, 3, 4, 5, and 6). Comparing PRO-seq and 3’-RNA-seq measurements (Supplementary Figure 7) revealed a moderate relationship (R^2^ values of 0.16, 0.35, 0.38 and 0.45 for the four species, respectively), consistent with transcript stability contributing strongly to transcript abundance in these four legume species.

### 4.2. Promoter proximal pausing is prevalent in invert repeat lacking legumes (IRLC)

While promoter proximal pausing appears to be a key transcriptional regulatory mechanism in animals, the role for this mechanism in plants is less clear. Land plants lack a homolog of negative elongation factor which plays a central role in promoter proximal pausing in metazoans [20]. Varying degrees of promoter proximal pausing have been reported across the limited number of species that have been investigated. In Arabidopsis, Hetzel et al [21] found a relative absence of pausing, while another researchers [22] found more pronounced pausing. Using PRO-seq, Lozano et al [23] found sharp promoter proximal pausing in maize and a more distributed but pronounce pausing profile in cassava. We found evidence of pausing in all four species with the overall profiles of faba bean, grass pea and barrel medic closely resembling cassava (Figure 4). The profile for pea was less pronounced. We also found enrichment of PRO-seq reads at the annotated gene ends in all species (Supplementary Figure 8). A depression in the signal directly overlapping the end of the annotated gene which was most pronounced in *Vicia faba*. As nearly all gene annotation for this species lack UTRs, this depression is centered at the end of the coding sequence.

**Figure 4.**
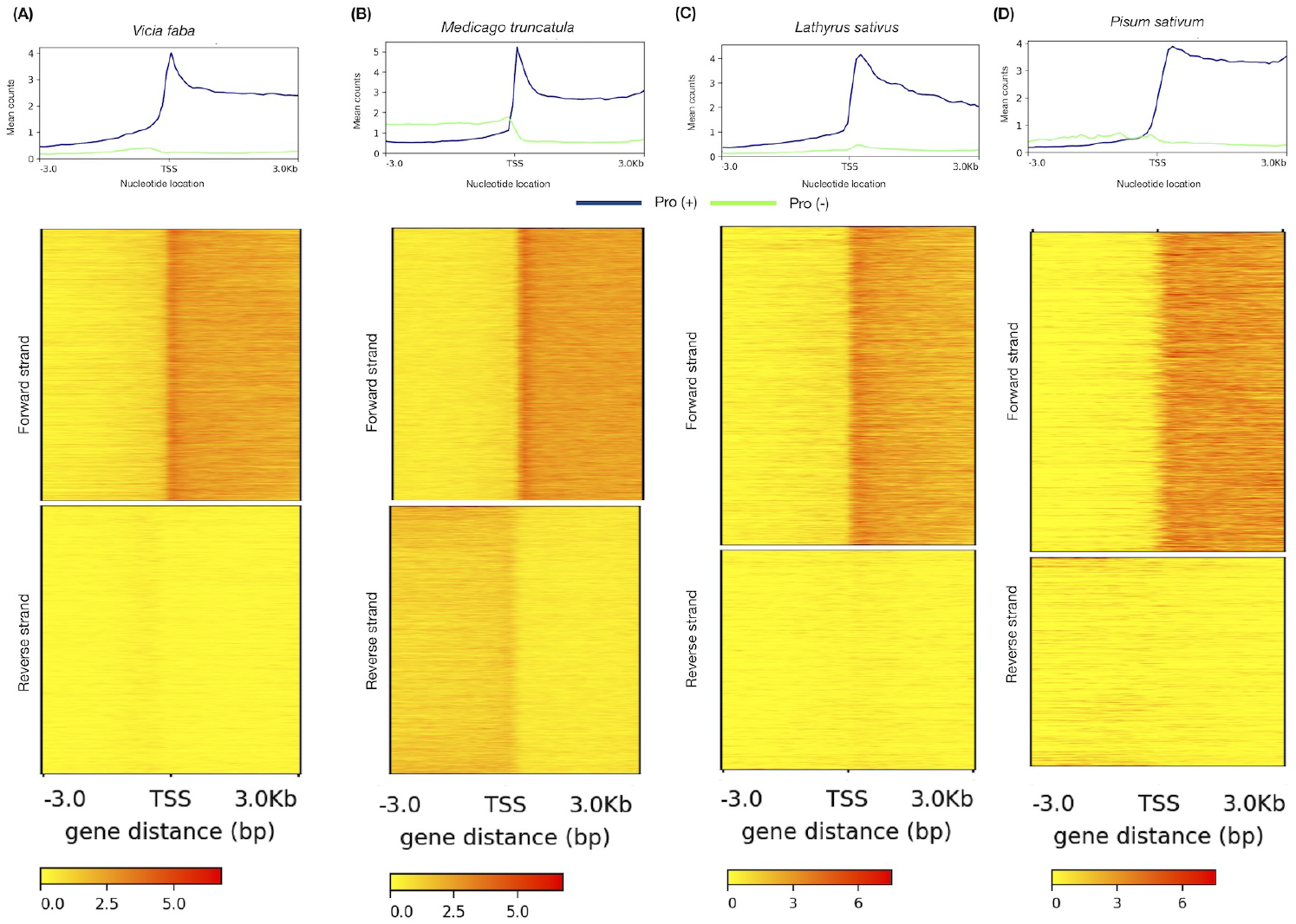
Accumulation of PRO-seq reads around the transcription start site (TSS) of four different plant species. Metaplot of PRO-seq signal from annotated genes normalized for reads per bp per gene in *Vicia faba* (A), *Medicago truncatula* (B), *Lathyrus sativa* (C) and *Pisum sativum* (D). Reads were aligned to the TSS and the TTS in both sense (blue) and antisense (green) directions relative to the direction of gene transcription.

### 4.3. PRO-seq measurements are more conserved across related species than 3-RNA-seq measurements

Washburn *et al*. [7] showed that interspecific comparisons of ortholog gene expression can provide an information-dense source of training data for gene expression modelling. We scaled up this approach using four closely-related legume species and included PRO-seq to demonstrate independent modelling of transcription. Single copy orthologs displayed a correlation at the level of transcription and transcript abundance, with a stronger correlation at the level of transcription. As shown in the Supplementary Figure 9, the box plot indicates the average R^2^ values of the ortholog species pairs in PRO-seq are generally higher than in 3-RNA-seq expression data. More detailed figures can be found in Supplementary Figure 10.

### 4.4. PlanTT predicts expression difference from 3’-RNA-seq data

We first evaluated PlanTT with the tower architecture introduced in Section 3.2 and compared it to the model of Washburn *et al*. [7] in the prediction of gene expression difference (i.e., *d*_*a,b*_). We implemented the binary classification model as described in [7] with Pytorch [24] 2.1.0. The model was trained used binary cross-entropy as the loss function and area under receiver operating characteristic (ROC) curve as the early stopping metric. Rather than encoding missing bases using an additional dimension, we soft encoded them as in PlanTT (see Section 3.2). Further, considering that Washburn’s model is not equivariant to sequence permutation by construction, we augmented its training sets by including flipped versions of the pairs of orthologs. This helps the model achieve permutation equivariance through training, therefore leveling its comparison with PlanTT. A flipped pair of orthologs is obtained by swapping the sequences and changing the sign of the targets (i.e., **s**_*a*_, **s**_*b*_, *d*_*a,b*_ → **s**_*b*_, **s**_*a*_, − *d*_*a,b*_).

Table 1 shows regression and classification metrics for both models. For the PlanTT model, gene pairs are recognized to be correctly classified if the sign of their predicted effect size matches the sign of the ground truth. No regression metrics are reported for the Washburn *et al*. classification model, which does not predict the magnitude of the expression difference.

**Table 1:** Comparison of PlanTT to previous results for predicting differential expression in the 3’-RNA-seq data.

| | Balanced accuracy | F1-score | Spearman R | $R^2$ |
| --- | --- | --- | --- | --- |
| Washburn [7] | 0.640 | 0.636 | - | - |
| PlanTT | 0.604 | 0.594 | 0.375 | 0.145 |

PlanTT is able to predict the magnitude of expression difference between unseen orthologs, with a Spearman rank correlation of 0.375 and an R^2^ of 0.145. The Washburn *et al*. model was able to outperform the proposed contrastive learning model in the binary classification metrics (balanced accuracy and F1 score). This is expected, as the previous model was trained on the binary classification task specifically. The PlanTT model was trained to estimate the magnitude of the expression difference, and is only able to estimate the binary characterization incidentally. Intuitively, for the many genes in the dataset with small ground-truth differences in expression, PlanTT will receive a low loss signal if the prediction is close to zero, but whether it is slightly negative or slightly positive is inconsequential.

### 4.5. Sequential Training Using 3’-RNA-seq and PRO-Seq

We explored the prospect of using one type of count data as a pre-training task, in order to potentially improve performance on the other dataset as a downstream task. Pre-training in this way is motivated by the fact that the measurements are inherently connected – the mRNA abundance of a gene is the product of its transcription rate and its stability. While stability is a key factor of a transcript’s life expectancy, transcription rate itself tells us about the quantity of messages sent by the cell to create a protein, thus possibly a good proxy of reading abundance. Table 2 shows performance using PRO-Seq and 3’-RNA-seq alone, as well as using each as the pre-training and downstream task, using the same cross-validation splits as in Section 4.4. For these experiments, no weights were frozen and training was started on the downstream task using the best weights obtained on the pre-training task as the initialization.

**Table 2:** Results of transfer learning with PlanTT.

|  | Balanced accuracy | F1-score | Spearman R | R<sup>2</sup> |
| --- | --- | --- | --- | --- |
| PRO-Seq | 0.532 | 0.532 | 0.099 | 0.001 |
| 3'-RNA-seq → PRO-Seq | 0.551 | 0.551 | 0.181 | 0.017 |
| 3'-RNA-seq | 0.604 | 0.594 | 0.375 | 0.145 |
| PRO-Seq → 3'-RNA-seq | 0.606 | 0.596 | 0.375 | 0.151 |

We observe that PlanTT can achieve results which are marginally better than random, with a balanced accuracy of 0.532, when estimating transcription rate differences between the orthologs. Further, we see that using the resulting weights of the latter experiment as a starting point for the downstream task leads to a slight increase in R^2^. We see a larger impact when PRO-Seq data is considered for the downstream task (3’-RNA-seq → PRO-Seq), although the overall performance on PRO-Seq remains limited. While using both datasets for training had a positive impact on both tasks, the limited impact of the PRO-seq suggests that this modelling approach is more amenable to modelling post-transcriptional processes such as transcript stability. This is not unexpected since the windows included here include the region expected to regulate stability (the transcribed region), but may miss important regulatory sequences located further away gene sequence that govern transcription.

### 4.6. Analysis of feature importance

To visualize how PlanTT models the interaction between nucleotide sequences and differences in gene expression, we conducted a feature importance analysis using the implementation of DeepLift-Rescale [25] provided by the *Captum* [26] library. In Figure 5, we present the average of the absolute values of the attribution scores for each nucleotide location and all of the sequences present in the test sets during the cross-validation executed in Section 4.4.

**Figure 5.**
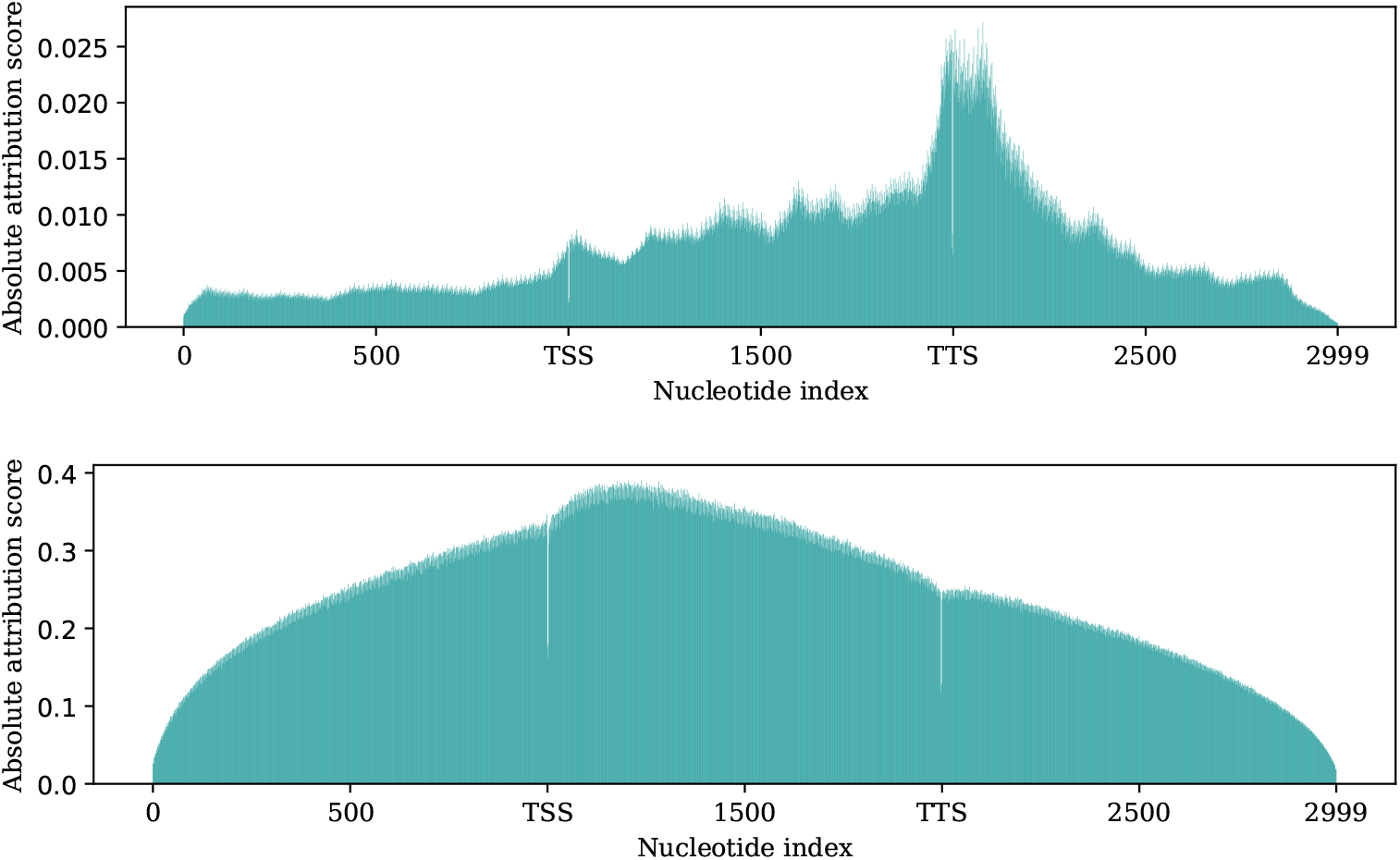
Average of the absolute value of the attribution scores for each nucleotide position using 3’-RNA-seq (top) and PRO-seq (bottom). A tensor full of zeros is used as the baseline for DeepLift. TSS: transcription start site; TTS: transcription termination site.

We observe that the predominant features are the bases surrounding the transcription termination site, which is similar to the findings of Washburn et al [7], although the Washburn model was trained with a classification goal and different species. This result also aligns with previous work highlighting the importance of the terminator region in regulating gene expression in plants [27]. While this finding supports the role for transcript stability in gene expression variation, the 3’-bias may also reflect the relative difficulty of learning transcript stability versus transcription.

### 4.7. Prediction of variant effects with PlanTT

PlanTT can be employed to predict the effect of single-base edits to a gene sequence, with the goal of increasing or decreasing its transcript abundance. Figure 6 demonstrates this for a random subset of sequences from *Vicia faba* not seen during training. Similar heatmaps for other species are shown in Supplementary Figures 12, 13, and 14.

In the context of gene editing, we can take advantage of the design of the head to return the expected changes in percentage rather than in absolute measures. Precisely, let **z** and **z**_*m*_ be the embeddings associated to a gene sequence and any of its mutations *m* respectively. According to Equation 1, it is possible to estimate the expression of the original sequence with sum(**z**) and the one of its mutation with sum(**z**_*m*_). Therefore, the percentage in change can be represented as:

**Figure 6.**
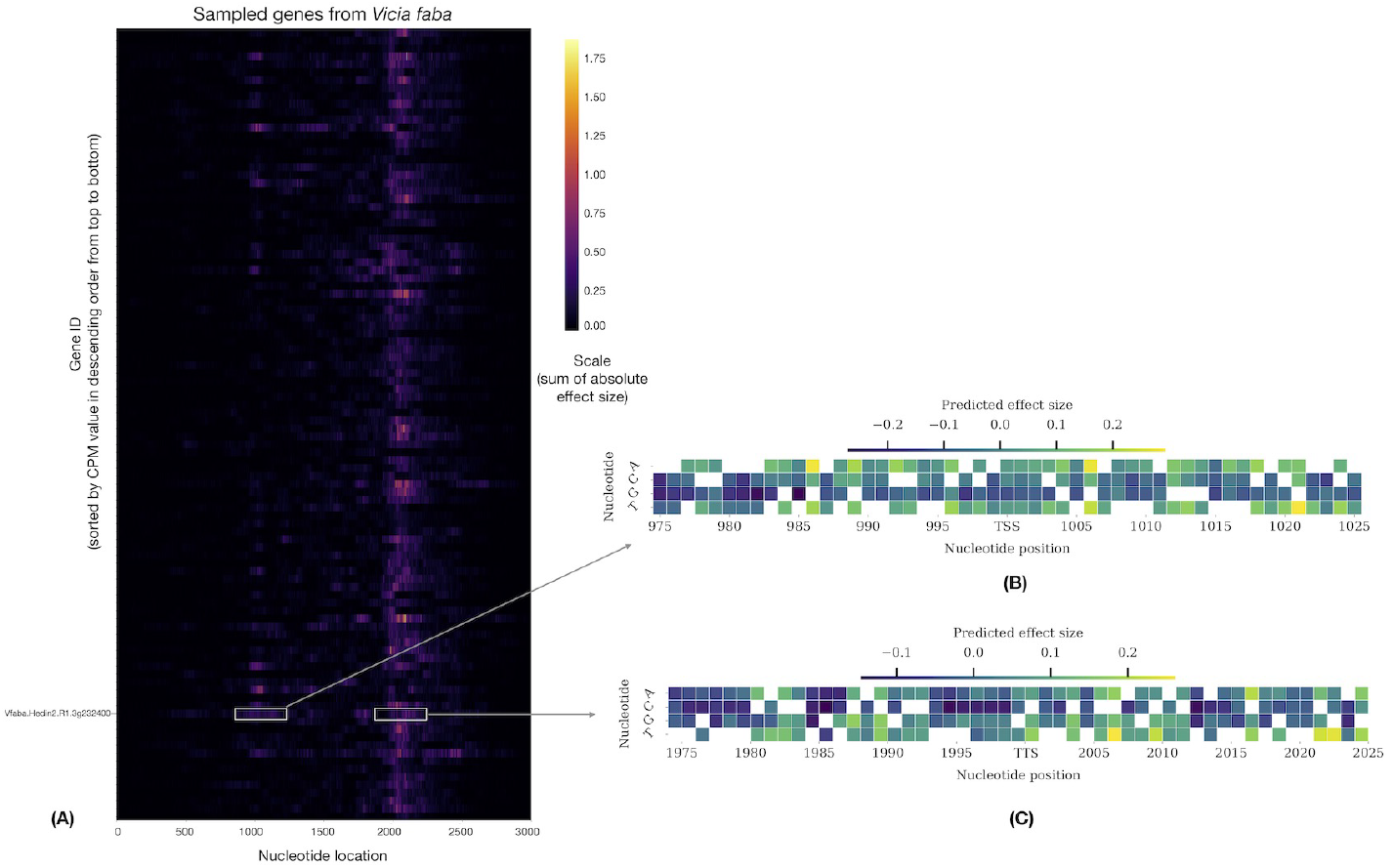
Left, heat map of predicted gene expression impact of base modifications for representative Vicia faba genes. Each row represents a single gene, at each base the edit with the greatest impact is plotted. The Transcript start and end sites are located at positions 1000 and 2000, respectively. Gene order is based on descending expression level. Downsampled to 100 genes. Right, prediction of gene expression impacts for a representative gene at the gene start and end. Original bases are left blank.

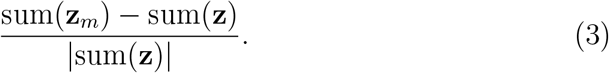

In Equation 3, it is necessary to take the absolute measure of the denominator as the sum of the elements in an embedding can be negative, meaning that the expression of a gene is predicted to be repressed. It is to be noted that we implemented our framework so that the denominator is replaced by a small value of *ϵ* if it is null. In our case, we set *ϵ* to 1*e −* 4.

To transmute a given sequence to a higher or lower expression level, we can apply an editing process in which we iteratively select a nucleotide position and select a modification that maximizes or minimizes the predicted expression of a sequence based on its latest state. Let **s**^(0)^ be a sequence in its original state, *M* be a maximal number of single-base modifications, and *ρ*(*s, i, j*) a function that replaces the *i*-th nucleotide of any sequence *s* by the value *j*, we can define the editing process as the creation of an ordered set of pairs 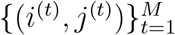 such that

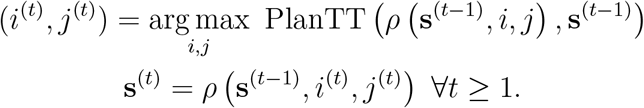

To ensure finding modifications that do not decrease transcript abundance, the procedure is stopped if the greatest prediction returned by PlanTT within an iteration is negative or equal to zero. As a hill climbing algorithm, the technique is not guaranteed to find the optimum ordered set of edits (i.e. the global maximum) [28], but provides a first approximation for sequence editing. An implementation of this algorithm is provided for reference.

## 5. Conclusion

In this work we described PlanTT, a neural network architecture trained via contrastive learning which is able to predict differential gene expression in ortholog pairs based on their promoter and terminator sequences. Unlike previous models which only predict which allele is more highly expressed, we have demonstrated that PlanTT is able to predict the difference in expression in terms of effect size. As a means to interpret and compare DNA sequences at the gene regulatory level, this type of model has potential for broad utility in plant biology and breeding, such as identifying functional variation, and informing gene editing strategies.

## Supporting information

NRC_2025_Manuscript_supplementary_files

## Availability and implementation

The implementation of PlanTT is available in the following GitHub repository: https://github.com/Amii-Open-Source/PlanTT. Please address the corresponding authors if you have questions of access.

