## Supplementary material for "Contrastive modelling of transcription and transcript abundance in legumes using PlanTT": NRC_2025_Manuscript_supplementary_files

### 1 Supplementary Figures

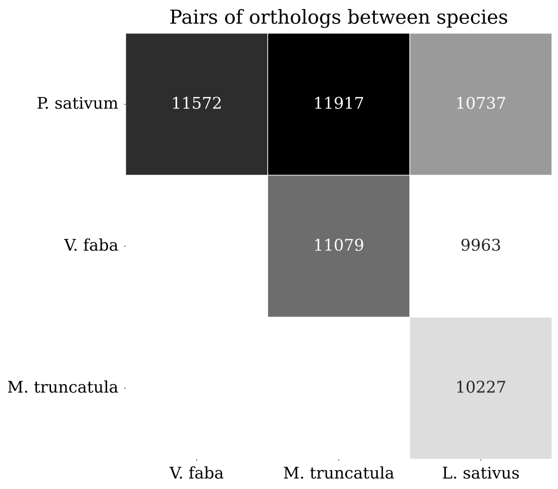


Supplementary Figure 1: Number of pairs of ortholog genes between each species.

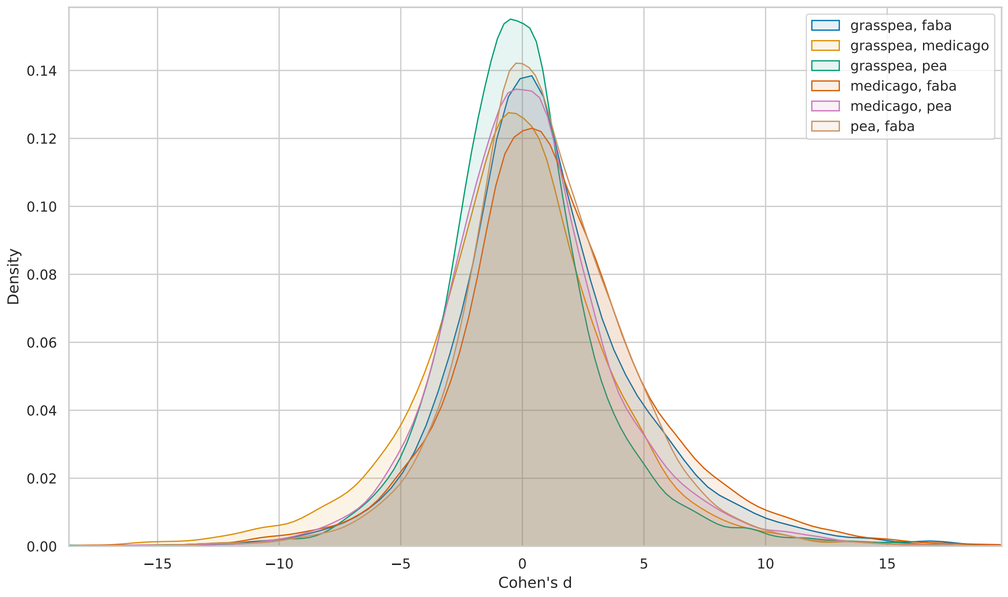


Supplementary Figure 2: The distribution of the effect size - Cohen's $d$ from each species pair. For each colored distribution plot, faba refers to *V. faba*; grasspea refers to *L. sativus*; medicago refers to *M. truncatula*; pea refers to *P. sativum*.

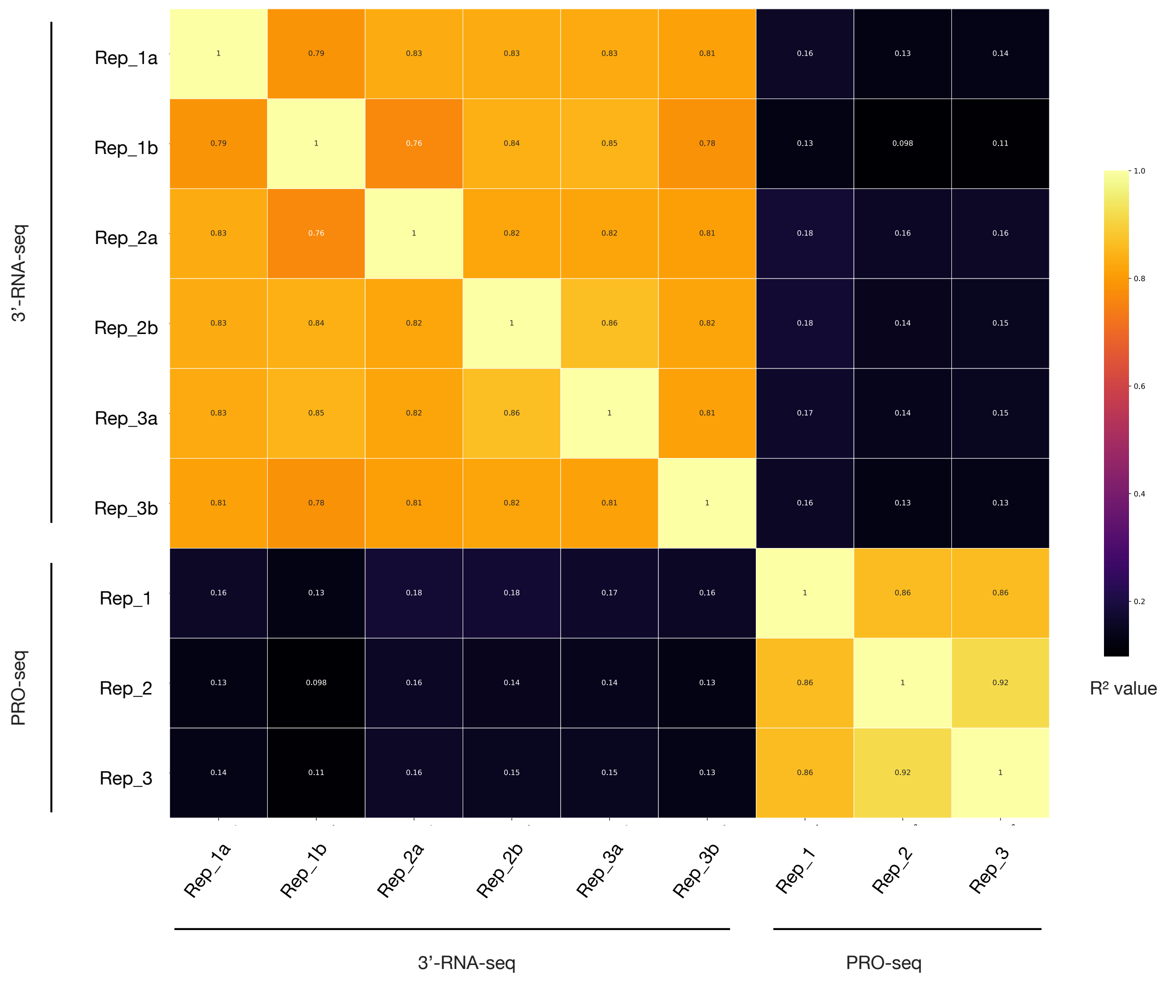


Supplementary Figure 3: The mean log abundance - counts per million (CPM) across replicates for the 3'-RNA-seq data compared to the standard deviation in *V. faba*. The scale shows the average Pearson $R^2$ value with p-value close to zero for each replicate comparison.

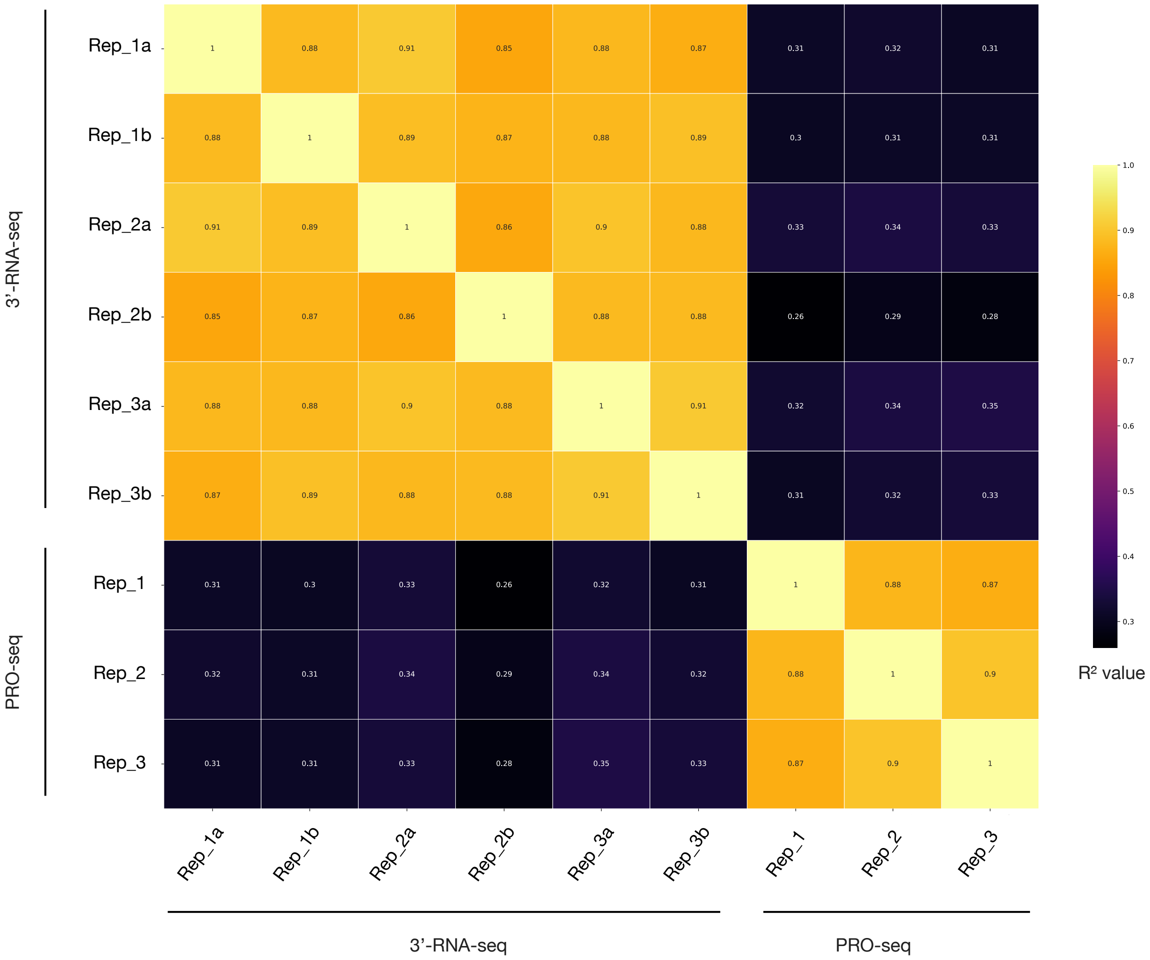


Supplementary Figure 4: The mean log abundance - counts per million (CPM) across replicates for the 3'-RNA-seq data compared to the standard deviation in *L. sativus*. The scale shows the average Pearson $R^2$ value with p-value close to zero for each replicate comparison.

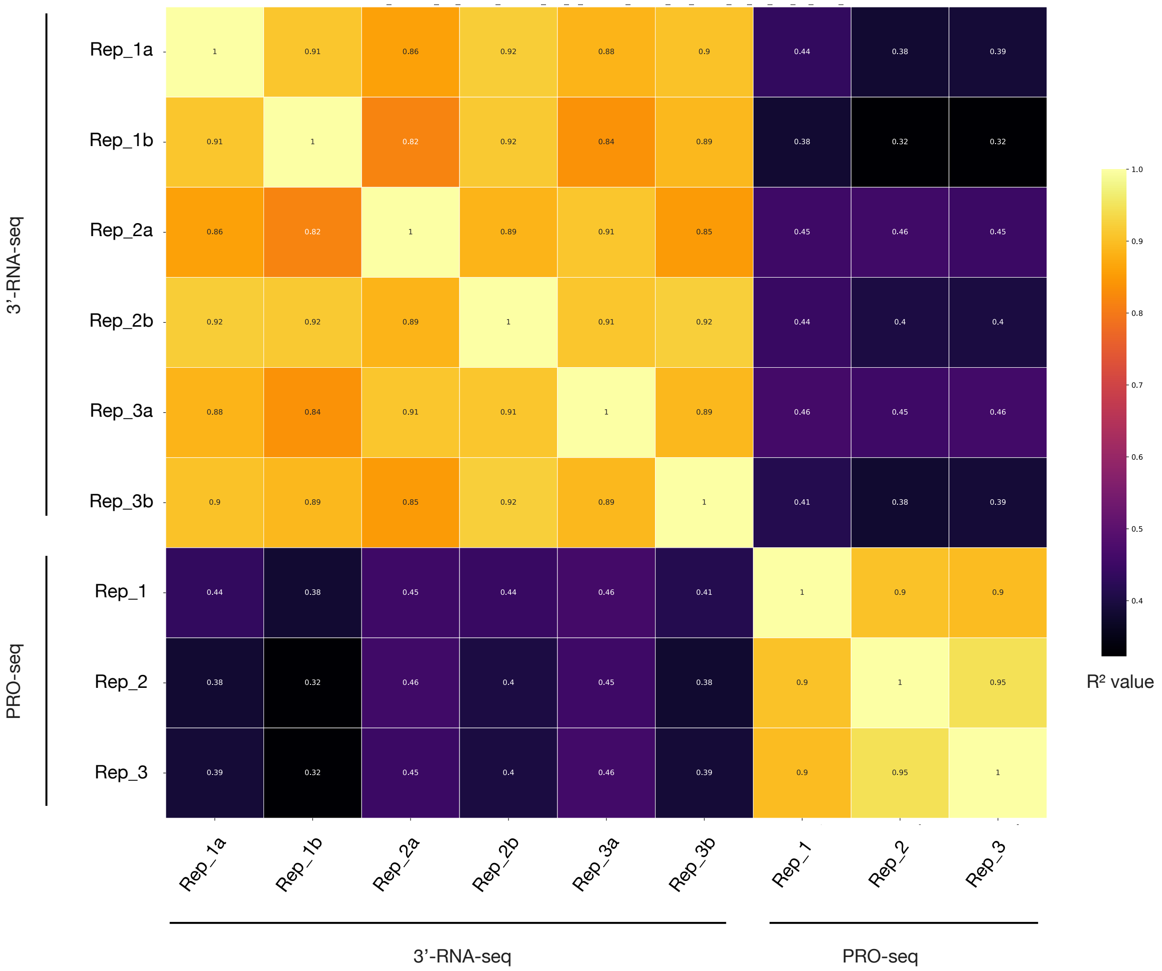


Supplementary Figure 5: The mean log abundance - counts per million (CPM) across replicates for the 3'-RNA-seq data compared to the standard deviation in *M. truncatula*. The scale shows the average Pearson $R^2$ value with p-value close to zero for each replicate comparison.

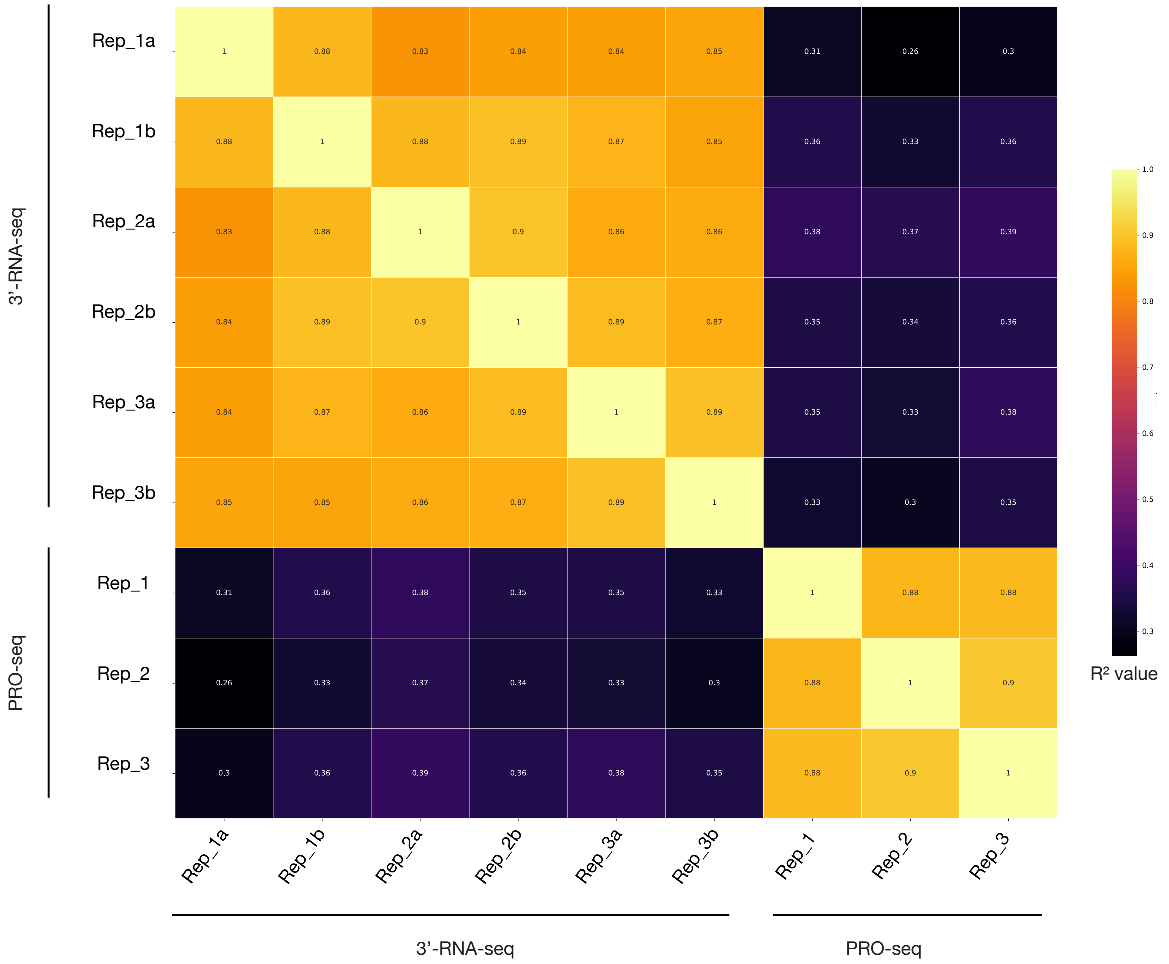


Supplementary Figure 6: The mean log abundance - counts per million (CPM) across replicates for the 3'-RNA-seq data compared to the standard deviation in *P. sativum*. The scale shows the average Pearson $R^2$ value with p-value close to zero for each replicate comparison.

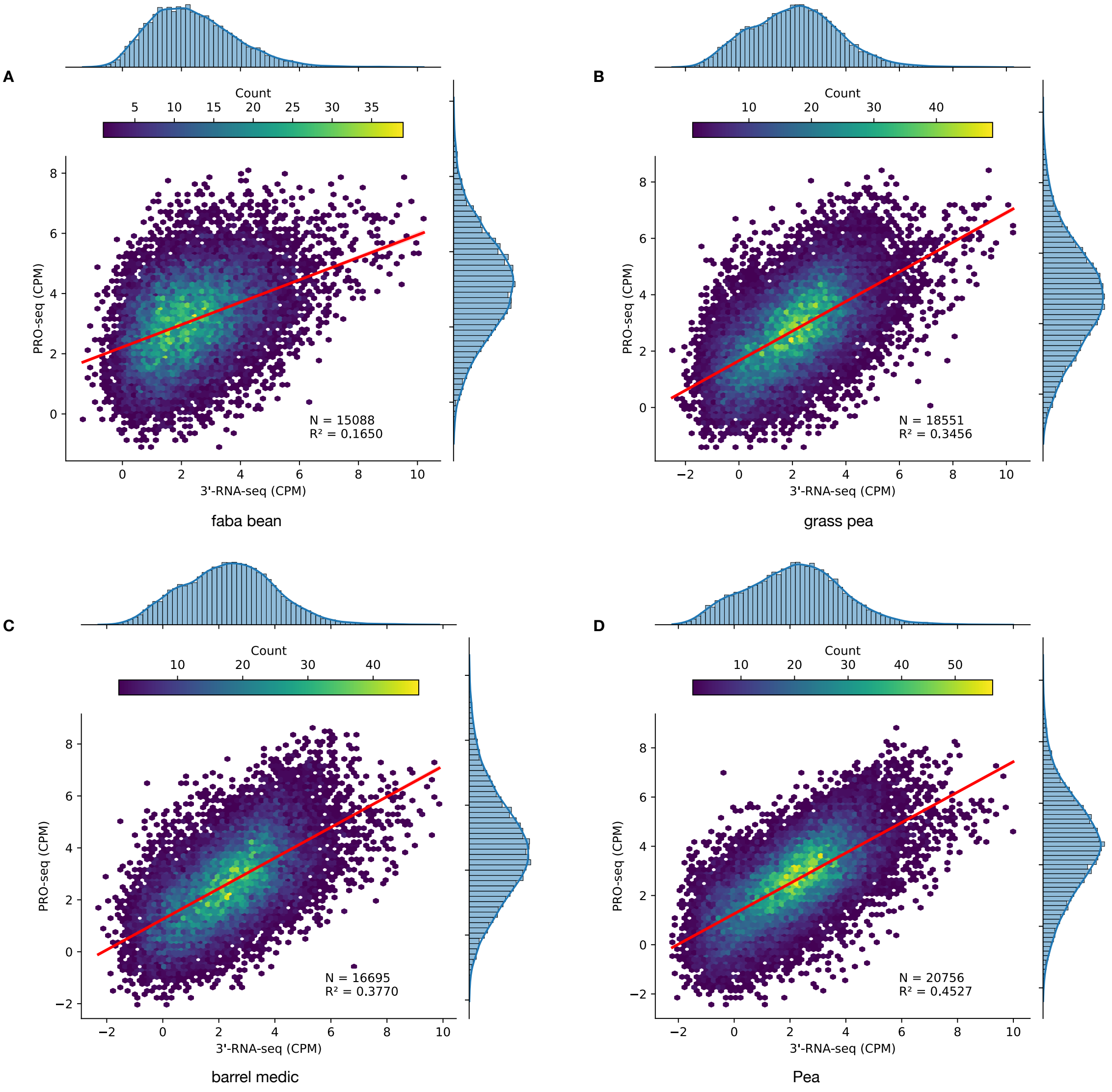


Supplementary Figure 7: Comparison of the mean log abundance - counts per million (CPM) for each species in the 3'-RNA-seq data and PRO-Seq data in Hexbin plot, organized by same gene. Unexpressed genes have been removed. For each plot, the number of genes (N) and the Pearson's coefficient ($R^2$) are shown. Faba bean refers to *V. faba*; Grass pea refers to *L. sativus*; Barrel medic refers to *M. truncatula*; Pea refers to *P. sativum*.

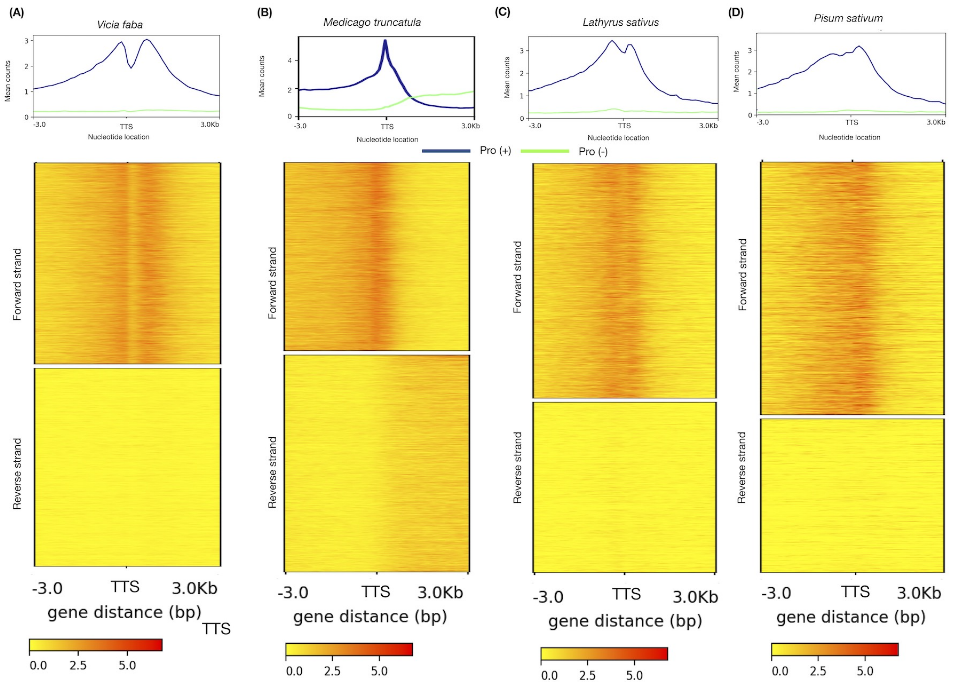


Supplementary Figure 8: Accumulation of PRO-seq reads around the transcription termination site (TTS) of four different plant species. Metaplot of PRO-seq signal from annotated genes normalized for reads per bp per gene in *Vicia faba* (A), *Medicago truncatula* (B), *Lathyrus sativa* (C) and *Pisum sativum* (D). Reads were aligned to the TSS and the TTS in both sense (blue) and antisense (green) directions relative to the direction of gene transcription.

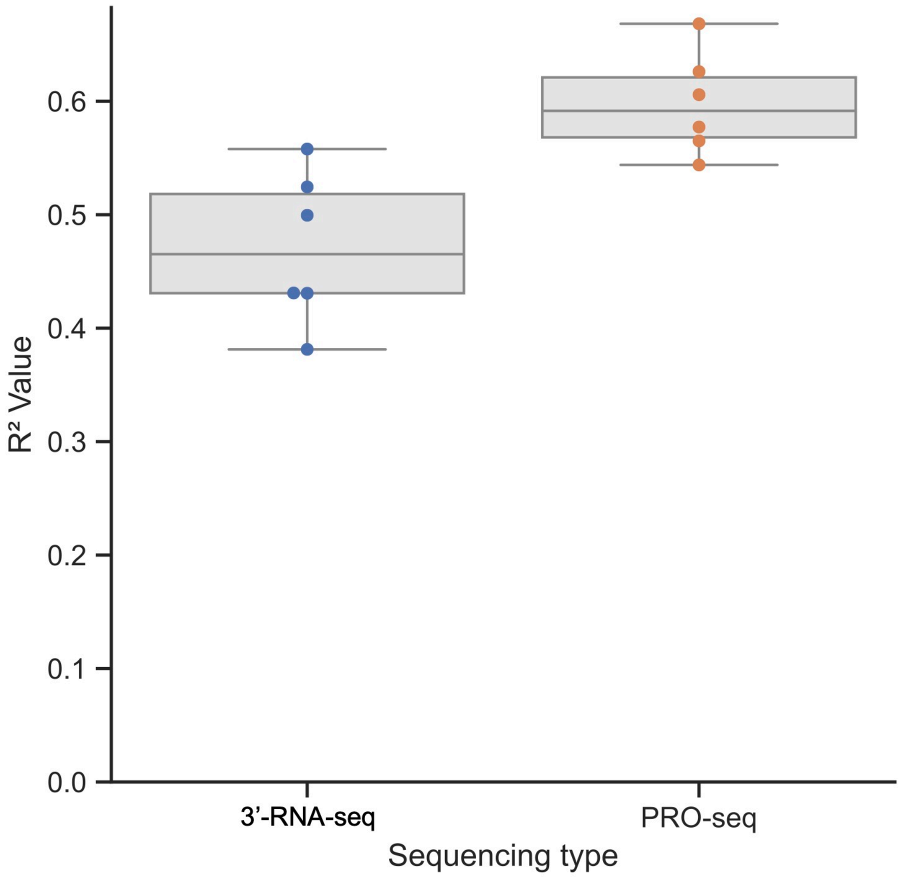


Supplementary Figure 9: Comparison of the mean log abundance - counts per million (CPM) in Pearson $R^2$ value for each ortholog in the 3'-RNA-seq and PRO-seq data in Box plot, organized by species pair. Unexpressed genes have been removed.

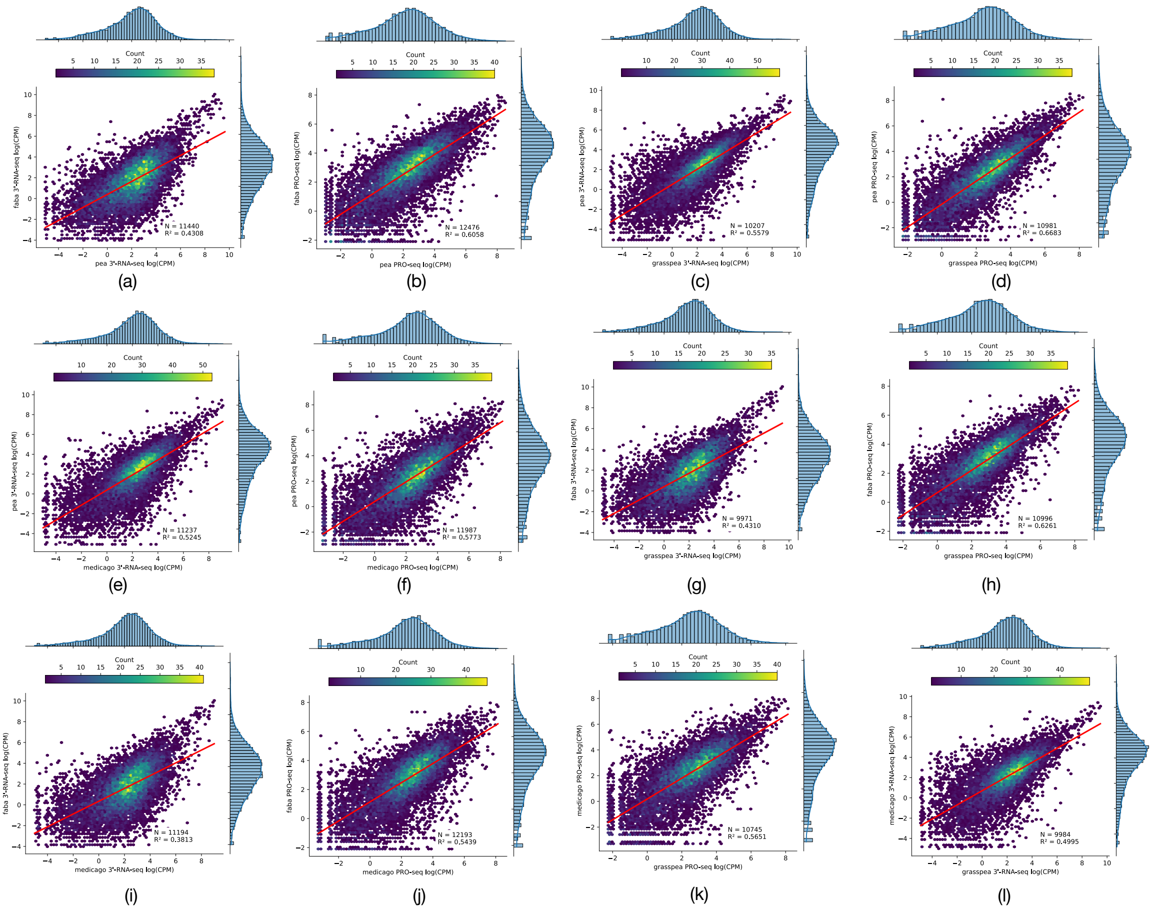


Supplementary Figure 10: Comparison of the mean log abundance - counts per million (CPM) for each ortholog in the 3'-RNA-seq data and PRO-Seq data in Hexbin plot, organized by species pair. Unexpressed genes have been removed. For each plot, the number of genes (N) and the Pearson's coefficient ($R^2$) are shown. Faba bean refers to *V. faba*; Grass pea refers to *L. sativus*; Barrel medic refers to *M. truncatula*; Pea refers to *P. sativum*

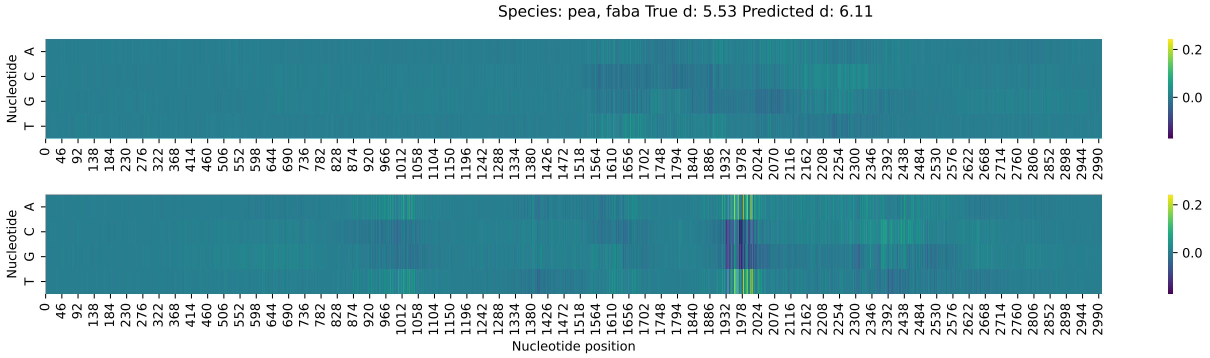


Supplementary Figure 11: Example of the model's predicted effect size (Cohen's $d$) change when making single-nucleotide edits to the *P. sativum* allele (top) and the *V. faba* allele (bottom) for a randomly selected gene. The color scale shows the predicted change in the effect size which results from changing the nucleotide.

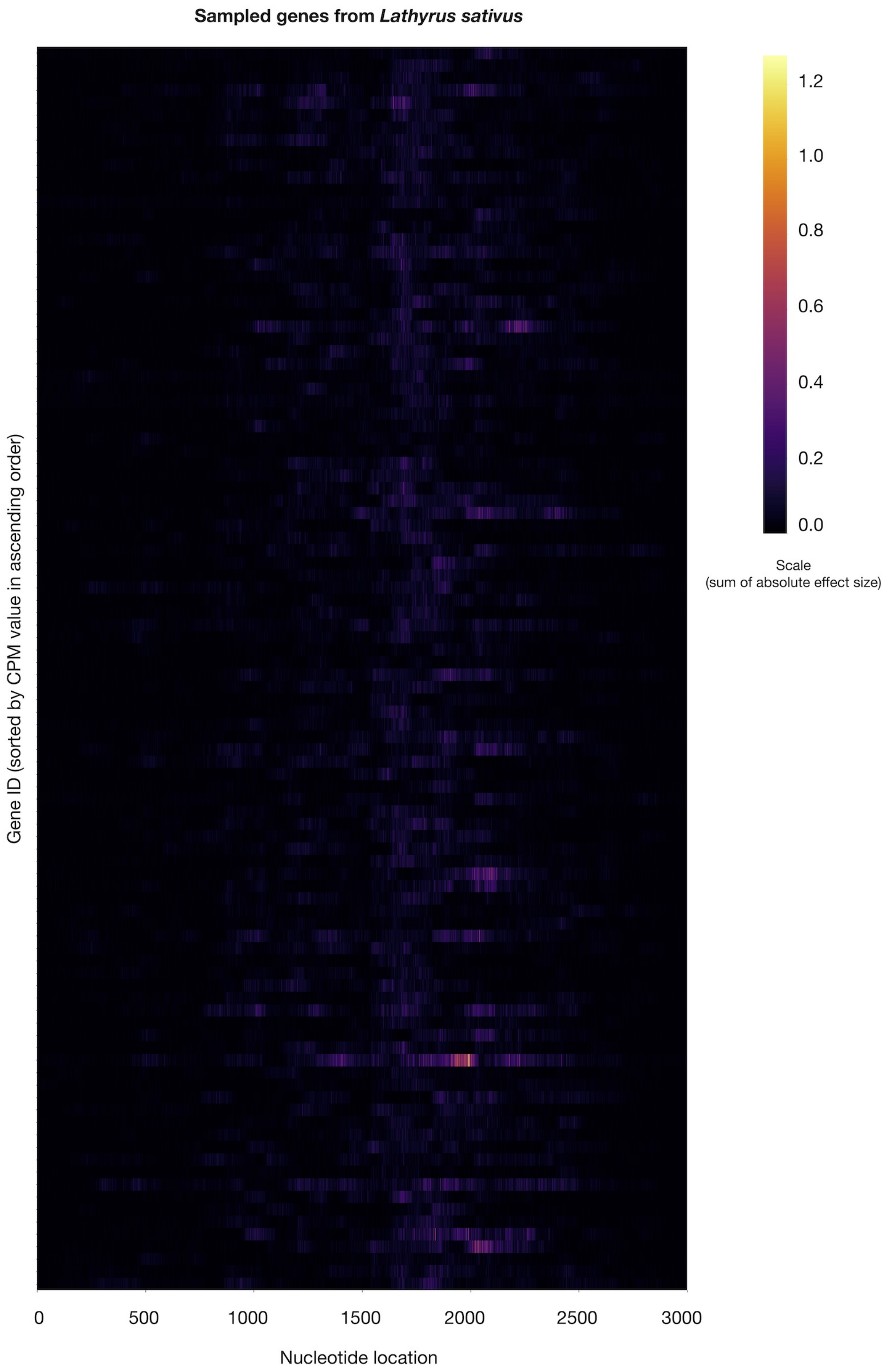


Supplementary Figure 12: Heatmap of predicted gene expression impact of base modifications for representative *L. sativus* genes. Each row represents a single gene, at each base the edit with the greatest impact is plotted. The Transcript start and end sites are located at positions 1000 and 2000, respectively. Gene order is based on descending expression level. Downsampled to 100 genes.

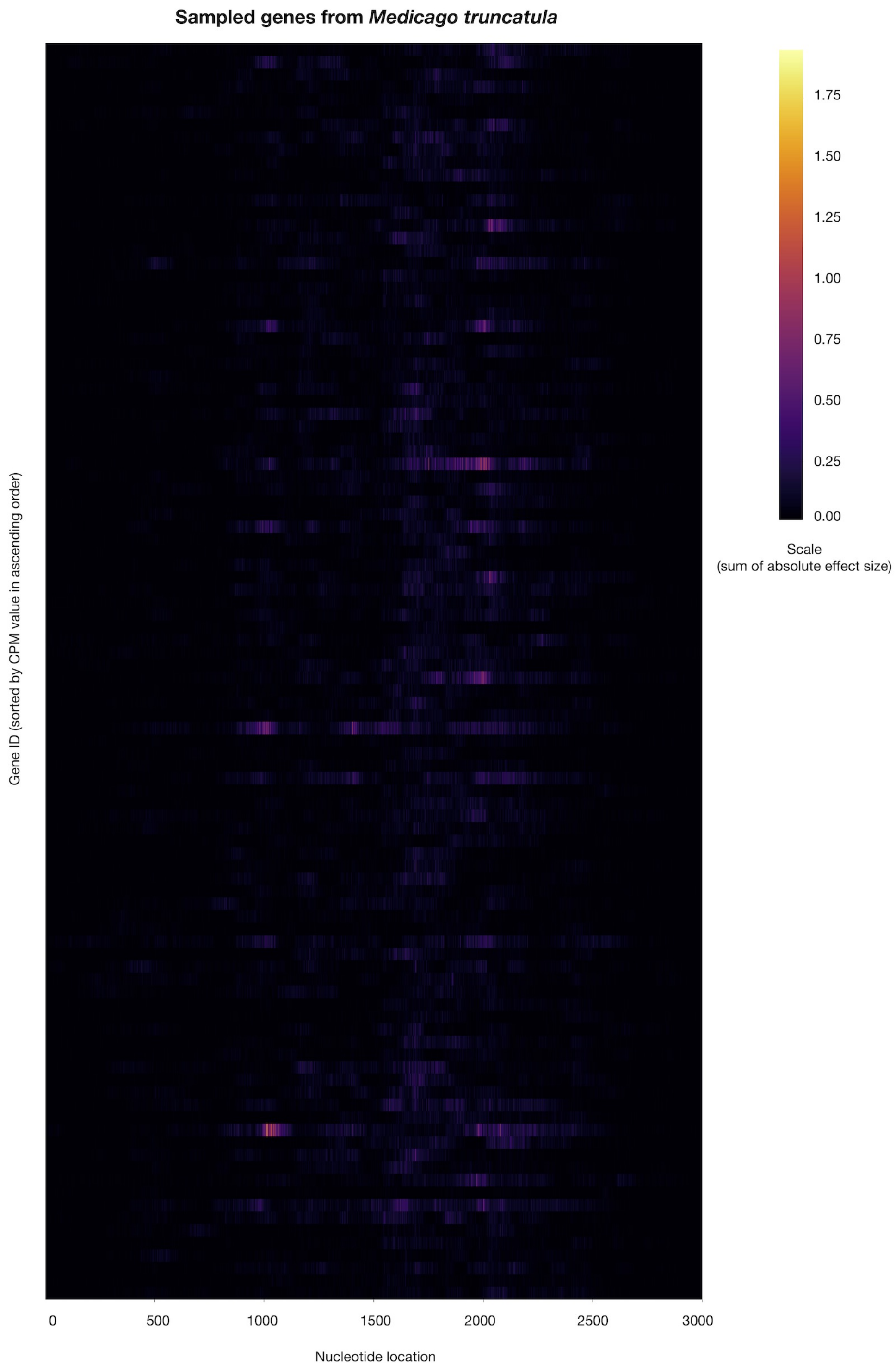


Supplementary Figure 13: Heatmap of predicted gene expression impact of base modifications for representative *M. truncatula* genes. Each row represents a single gene, at each base the edit with the greatest impact is plotted. The Transcript start and end sites are located at positions 1000 and 2000, respectively. Gene order is based on descending expression level. Downsampled to 100 genes.

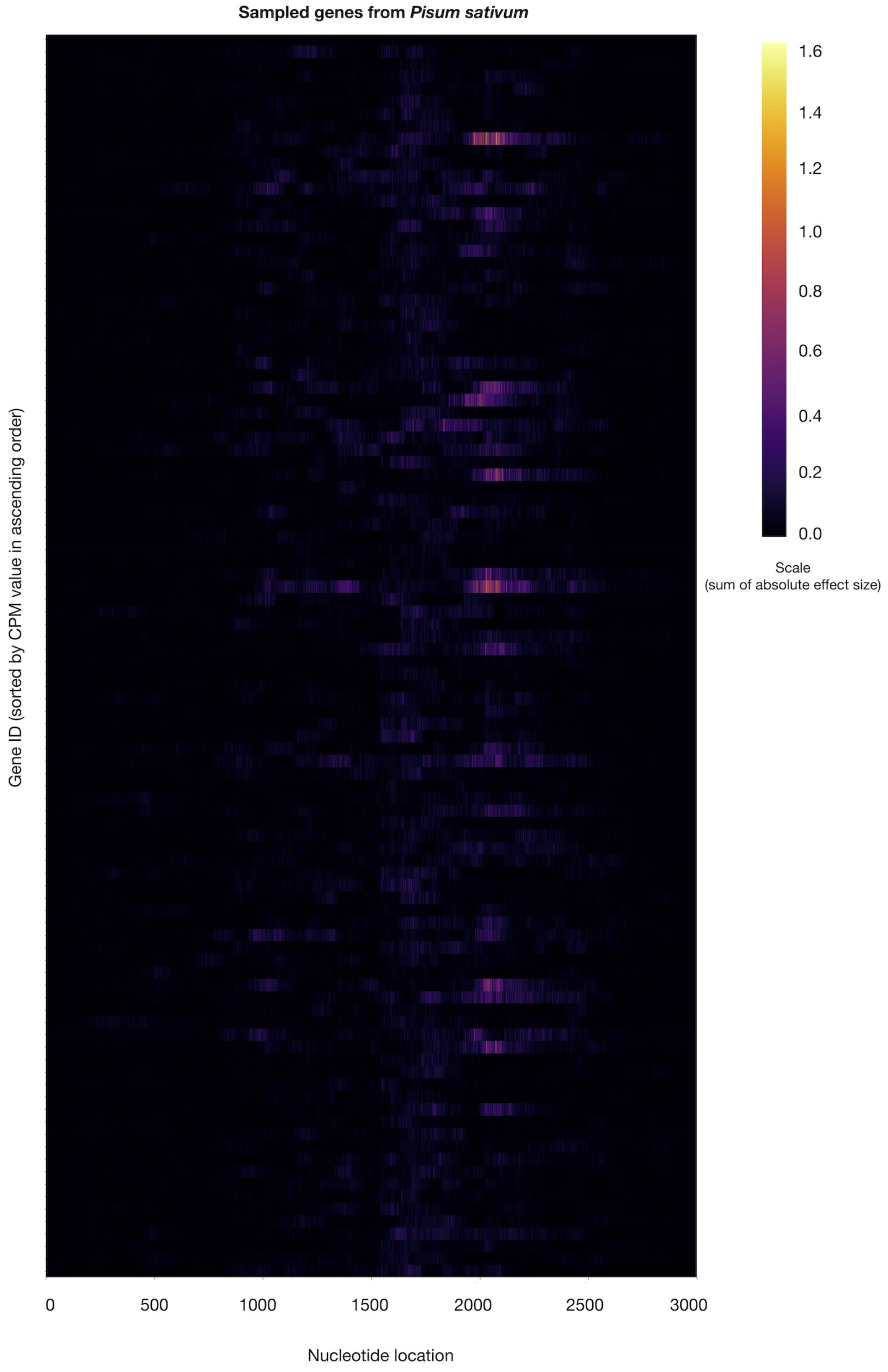


Supplementary Figure 14: Heatmap of predicted gene expression impact of base modifications for representative *P. sativum* genes. Each row represents a single gene, at each base the edit with the greatest impact is plotted. The Transcript start and end sites are located at positions 1000 and 2000, respectively. Gene order is based on descending expression level. Downsampled to 100 genes.

#### 2 Supplementary Proofs and Theorems

**Theorem 1** *Let* $f : (\mathbb{R}^{D\times L}, \mathbb{R}^{D\times L}) \rightarrow \mathbb{R}$ *be a model following PlanTT architecture, with* $t : \mathbb{R}^{D\times L} \rightarrow \mathbb{R}^{D'\times 1}$ *being its shared tower and* $h : \mathbb{R}^{D'\times 1} \rightarrow \mathbb{R}$ *being its head, such that*

$$f(\mathbf{s}_a, \mathbf{s}_b) = h\left(t(\mathbf{s}_a) - t(\mathbf{s}_b)\right).$$

*The function* $f$ *is a valid model of gene expression difference (i.e., satisfies Definition 1) if* $h$ *is an odd function, meaning that* $h(-\mathbf{x}) = -h(\mathbf{x})$ *for any* $\mathbf{x} \in \mathbb{R}^{D'\times 1}$*.*

**Proof 1** *Let* $f : (\mathbb{R}^{D\times L}, \mathbb{R}^{D\times L}) \rightarrow \mathbb{R}$ *be a model following PlanTT architecture, with* $t : \mathbb{R}^{D\times L} \rightarrow \mathbb{R}^{D'\times 1}$ *being its shared tower and* $h : \mathbb{R}^{D'\times 1} \rightarrow \mathbb{R}$*, an odd function, being its head. Considering these assumptions, we have that:*

$$\begin{aligned} f(\mathbf{s}_a, \mathbf{s}_b) &= h\left(t(\mathbf{s}_a) - t(\mathbf{s}_b)\right) \\ &= h\left(-(t(\mathbf{s}_b) - t(\mathbf{s}_a))\right) \\ &= -h\left(t(\mathbf{s}_b) - t(\mathbf{s}_a)\right) \\ &= -f(\mathbf{s}_b, \mathbf{s}_a), \end{aligned}$$

*for any* $\mathbf{s}_a, \mathbf{s}_b \in \mathbb{R}^{D'\times 1}$*. From that result, we can further see that:*

$$\begin{gathered} f(\mathbf{s}_a, \mathbf{s}_b) = -f(\mathbf{s}_b, \mathbf{s}_a) \\ \implies f(\mathbf{s}_a, \mathbf{s}_a) = -f(\mathbf{s}_a, \mathbf{s}_a) \\ \implies 2f(\mathbf{s}_a, \mathbf{s}_a) = 0 \\ \implies f(\mathbf{s}_a, \mathbf{s}_a) = 0 \end{gathered}$$

**Proof 2** *Let* $\mathbf{x} \in \mathbb{R}^{D'\times 1}$ *be a vector for which each element is denoted as* $x_i$*, with* $i = 1, \ldots, D'$*. Let also* $h : \mathbb{R}^{D'\times 1} \rightarrow \mathbb{R}$ *be defined as the sum of the elements contained in any vector within* $\mathbb{R}^{D'\times 1}$*, such that:*

$$h(\mathbf{x}) = \sum_{i=1}^{D'} x_i.$$

*We observe that:*

$$h(-\mathbf{x}) = \sum_{i=1}^{D'} -x_i = -\sum_{i=1}^{D'} x_i = -h(\mathbf{x})$$
